# MultiGAI: Global Attention-Based Integration of Single-Cell Multi-Omics with Batch Effects Correction

**DOI:** 10.64898/2025.12.18.692200

**Authors:** Jinyu Zhang

**Affiliations:** Xiamen University; National Institute for Data Science in Health and Medicine, Xiamen University

**Keywords:** single-cell multi-omics integration, spatial transcriptomics, batch effects correction, global attention

## Abstract

Single-cell multi-omics technologies allow the simultaneous measurement of multiple molecular modalities within the same cell, such as gene expression and chromatin accessibility (scRNA-seq + scATAC-seq) or gene expression and cell surface protein abundance (scRNA-seq + ADT), providing a multidimensional perspective on cellular states and regulatory mechanisms. However, these modalities often differ substantially in their distributions and noise levels and are affected by technical biases during experimental batches and sample processing, which can obscure true biological signals. To address these challenges, we present MultiGAI, a variational autoencoder (VAE) framework with a global attention mechanism. MultiGAI integrates global information from the dataset during encoding and employs specially designed components to limit the propagation of batch information. This design allows effective batch effects correction while preserving key biological signals, generating high-quality latent representations of cells. Notably, in addition to performing well on single-cell multi-omics data, MultiGAI also demonstrates good performance in batch effects correction for single-cell transcriptomic data and in the integration of spatial transcriptomic data. Overall, MultiGAI provides a novel strategy for batch effects correction while retaining biological information, offering new insights for future single-cell multi-omics data integration.

## 1 Introduction

The development of single-cell multi-omics technologies has significantly advanced our understanding of cellular states and their regulatory mechanisms. Compared with traditional single-omics sequencing, these technologies enable the simultaneous capture of multiple layers of molecular information within the same cell, such as scRNA-seq combined with scATAC-seq (e.g., 10x Multiome [1]) or scRNA-seq combined with ADT (e.g., CITE-seq [2]), providing complementary perspectives on cellular function and identity. Leveraging these data, researchers can not only dissect the molecular characteristics of different cell types within heterogeneous tissues but also further uncover cross-modality regulatory mechanisms and explore dynamic cellular changes in physiological and pathological processes [3, 4].

Nevertheless, integrating single-cell multi-omics data remains challenging. On one hand, different modalities exhibit substantial differences in dimensionality, noise characteristics, and measurement ranges; on the other hand, technical biases arising from experimental preparation, sequencing platforms, and batch effects can obscure true biological signals and compromise the accuracy of downstream analyses [3]. Therefore, effectively correcting for batch effects while obtaining high-quality low-dimensional representations of cells during integration remains a critical problem.

To address this challenge, researchers have proposed various batch effects correction strategies to mitigate the interference of batch information in single-cell multi-omics integration. These strategies can be analogized to the structure of a variational autoencoder (VAE) [5]: depending on where the correction occurs, methods can be categorized into three types—performing batch effects correction in the encoder, the latent space, or the decoder. Among these, latent space and decoder-based corrections are the most commonly used. For example, Seurat [6] employs latent space batch effects correction by identifying biologically similar cell pairs (anchors) from different batches in a low-dimensional space and then mapping cells from each batch to a shared space to correct batch effects. In contrast, MultiVI [7] and Multigrate [8] perform decoder-based batch effects correction by injecting batch information into the decoder and applying constraints through the loss function. A recent method, MIMA [9], represents an encoder-based batch effects correction strategy: it partitions the latent space into three subspaces—shared, modality-specific, and batch—achieving batch effects correction by disentangling the latent representations.

However, these methods still have certain limitations. For latent space batch effects correction, since no additional design is applied during dimensionality reduction, batch information may become entangled with biological signals in the latent space, making it difficult to fully preserve biological information during correction. For decoder-based batch effects correction, which is commonly used in current deep learning approaches, batch information contributes to reconstructing the observed data. Consequently, under the guidance of the reconstruction loss, the latent space often retains partial batch information, which may weaken the effectiveness of batch effects correction. In contrast, encoder-based batch effects correction has greater potential. Although information loss is inevitable during dimensionality reduction, if such loss is primarily limited to batch information, the latent space can retain more biological signals, resulting in higher-quality latent representations. Nevertheless, MIMA [9], despite being an encoder-based batch effects correction, lacks additional design in its encoder, allowing biological signals to leak into the batch space and leading to information loss. Moreover, non-deep learning methods generally focus on relationships between samples and the global structure of the dataset, thereby preserving overall information that is crucial for improving single-cell multi-omics integration. In contrast, existing deep learning integration methods usually process each sample independently and rarely exploit inter-sample relationships or global dataset structure.

In summary, to address the limitations of existing batch effects correction strategies, we propose an encoder-decoder joint batch effects correction strategy. Building on the original decoder-based batch effects correction, this strategy introduces a module in the encoder to suppress the flow of batch information while preserving the global information of the dataset, thereby more effectively retaining biological signals in the latent space.

Based on this strategy, we propose MultiGAI. Its core innovation lies in the encoder: by employing a global attention mechanism, the encoder can capture global information during dimensionality reduction and leverage this structure to suppress the flow of batch information, thereby enabling the latent space to better preserve biological signals. In addition, the model incorporates mechanisms to account for technical noise, such as sequencing depth [10], mitigating the impact of non-biological factors on data integration.

Experimental results demonstrate that MultiGAI achieves a balanced performance between batch effects correction and the preservation of biological information in single-cell multi-omics data integration. Moreover, it also performs well in batch effects correction for single-cell transcriptomics and in the integration of spatial transcriptomics data.

## 2 Methods

### 2.1 Global Attention Mechanism

Currently, the attention mechanism in Transformers [11] is widely used in tasks that require understanding global relationships due to its strong capability for contextual modeling. This mechanism dynamically weights information through interactions among queries (**Q**), keys (**K**), and values (**V**), thereby enhancing the model’s expressive power. In single-cell multi-omics data, we aim for the model to capture the contextual information of each cell. However, if all cells are treated as the key-value set {**K, V**}, both computational complexity and memory usage increase dramatically with the number of cells, making training difficult.

To address this issue, our method draws on the core idea of Perceiver [12] by introducing a global, trainable, and fixed-size key-value set {**K, V**}, which serves as a compressed representation of the entire dataset.

From a mechanistic perspective, in conventional attention mechanisms, the query vector **q** of each token interacts with the key-value set {**K, V**} to generate the representation of that token. Here, the key-value set is composed of the key vectors **k** and value vectors **v** of all tokens. However, in biological contexts, cells exhibit significant correlations. Therefore, a fixed-size key-value set {**K, V**} can serve as a compact approximation of the full key-value set containing all cells.

This architecture fundamentally constrains the expressive capacity of the network by limiting the available representation space. During training, batch information is explicitly injected into the decoder, whereas biological information can only be provided by the input data. Therefore, under the guidance of the reconstruction loss and the constraint of limited representational capacity, the model tends to prioritize capturing biological information in the latent space that is essential for reconstruction and can only be obtained from the input data, rather than batch information that is also accessible to the decoder. This mechanism forms an information bottleneck [13, 14], effectively suppressing the propagation of batch information.

From a biological perspective, we assume that the state of each cell can be represented as a combination of several fundamental cellular states. Our goal is to enable the key–value set {**K, V**} to learn these shared fundamental cellular states and to construct latent representations of cells accordingly, thereby obtaining a latent space with biological interpretability.

Specifically, MultiGAI learns a query vector **q**^(*m*)^ for the input data **x**^(*m*)^ of each modality *m*, which interacts with the key-value set of that modality via attention to produce a modality-specific intermediate representation **e**^(*m*)^. Subsequently, the intermediate representations of all modalities are averaged element-wise to obtain a joint query vector **q**. This joint query vector then interacts with the joint key-value set through attention to generate the joint representation **e**, which is further used to parameterize the distribution of the joint latent variable **z** via separate neural networks for its mean ***µ*** and standard deviation ***σ***.

The advantages of this method include:

1. **1.Efficient global context integration** The fixed global key-value set serves as a shared memory for the dataset, allowing each sample’s query vector **q** to access a set of condensed cross-sample features. This effectively enhances the cross-sample consistency and generalization of the latent representations, while avoiding the high computational cost of conventional attention mechanisms on large datasets.
2. **Dynamic information selection and information bottleneck** Through the interaction between the query vector **q** and the global key set {**K**}, the model can dynamically weight and extract the most relevant information from the global value set {**V**} for the current sample. Meanwhile, the global attention mechanism forms an information bottleneck [13, 14] during dimensionality reduction, effectively suppressing irrelevant batch information. As a result, the latent variable **z** flexibly integrates global information while retaining richer biologically relevant signals, thereby improving reconstruction quality and downstream analysis performance.

### 2.2 Model Architecture

MultiGAI is a cell data integration method based on the variational autoencoder (VAE) framework [5], which incorporates a global attention mechanism and employs a zero-inflated negative binomial (ZINB) distribution [10] as the observation model.

The overall workflow for processing single-cell transcriptomic data is as follows. MultiGAI first learns a query vector **q** from the single-cell transcriptomic data **x** and performs attention interactions with the key– value set {**K, V**} [11] to generate an intermediate representation **e**. Based on this intermediate representation, independent neural networks learn the parameters of the latent variable distribution (mean ***µ*** and standard deviation ***σ***), and latent variables **z** are sampled from this distribution using the reparameterization trick [5]. Finally, the latent variables **z** are concatenated with the one-hot encoded labels of the corresponding batch and fed into the decoder to generate the parameters of a ZINB distribution [10], which are then used to reconstruct the original data while accounting for sequencing depth. The model is trained by maximizing the log-likelihood of the observations and simultaneously minimizing the KL divergence between the latent variable distribution and the prior.

The overall workflow for processing single-cell multi-omics data is as follows: MultiGAI first learns the corresponding query vector **q**^(*m*)^ from the single-cell data **x**^(*m*)^ of each modality, and interacts with the key-value pairs {**K**^(*m*)^, **V**^(*m*)^} of that modality through attention [11] to generate the intermediate representation **e**^(*m*)^ for the modality. Then, the intermediate representations of all modalities are averaged element-wise to obtain a joint query vector **q**, which interacts with the joint key-value pairs {**K, V**} through attention to produce the joint representation **e**. Based on this unified representation, independent neural networks learn the parameters of the latent variable distribution (mean ***µ*** and standard deviation ***σ***), and sample the latent variable **z** from this distribution using the reparameterization technique [5]. Finally, the latent variable **z** is concatenated with the one-hot encoded labels of the corresponding batch and fed into the decoder to generate the ZINB distribution parameters [10] for each modality, which are then used to reconstruct the original data by incorporating the sequencing depth or total counts of each modality. The model is trained end-to-end by maximizing the log-likelihood of the observed data for each modality while minimizing the KL divergence between the latent variable distribution and the prior distribution. Additionally, to enhance the consistency between the intermediate representations of different modalities, a modality consistency constraint based on cosine similarity is introduced to further improve the representation of the joint latent variables.

For spatial transcriptomic data, although gene expression is essential, spatial information is also critical for understanding cellular states [15]. Therefore, we aim to obtain latent representations that incorporate spatial information and perform downstream tasks based on these representations, such as reconstructing gene expression. In MultiGAI, gene expression data and spatial information (e.g., slides and spatial coordinates) are treated as two separate modalities for integration. For the gene expression modality, the processing follows the same procedure as for single-cell multi-omics data. For the spatial modality, slides are first one-hot encoded (if data from multiple slides are integrated, each slide is one-hot encoded; if only a single slide is present, only the spatial coordinates are modeled). The spatial coordinates are then normalized and encoded using Fourier feature transformation [16]. The one-hot encoded slides and Fourier-encoded spatial coordinates are concatenated to form the input for the spatial modality. A neural network is then used to learn this spatial modality data, producing an intermediate representation **e**^(*m*)^, after which subsequent operations follow the same procedure as for single-cell multi-omics data. Notably, the global attention mechanism is not applied to the spatial modality, as spatial information is inherently independent of batch effects and represents objective records. In the output stage for the spatial modality, the model predicts the normalized spatial coordinates, and the mean squared error (MSE) is used as the loss function. All other processing steps are consistent with those used for single-cell multi-omics data.

Figure 1 illustrates the overall architecture of the MultiGAI model. Detailed mathematical formulations of the model, including the encoder, latent variable sampling, decoder, ZINB likelihood, and loss function, are provided in Appendix A.

**Figure 1.**
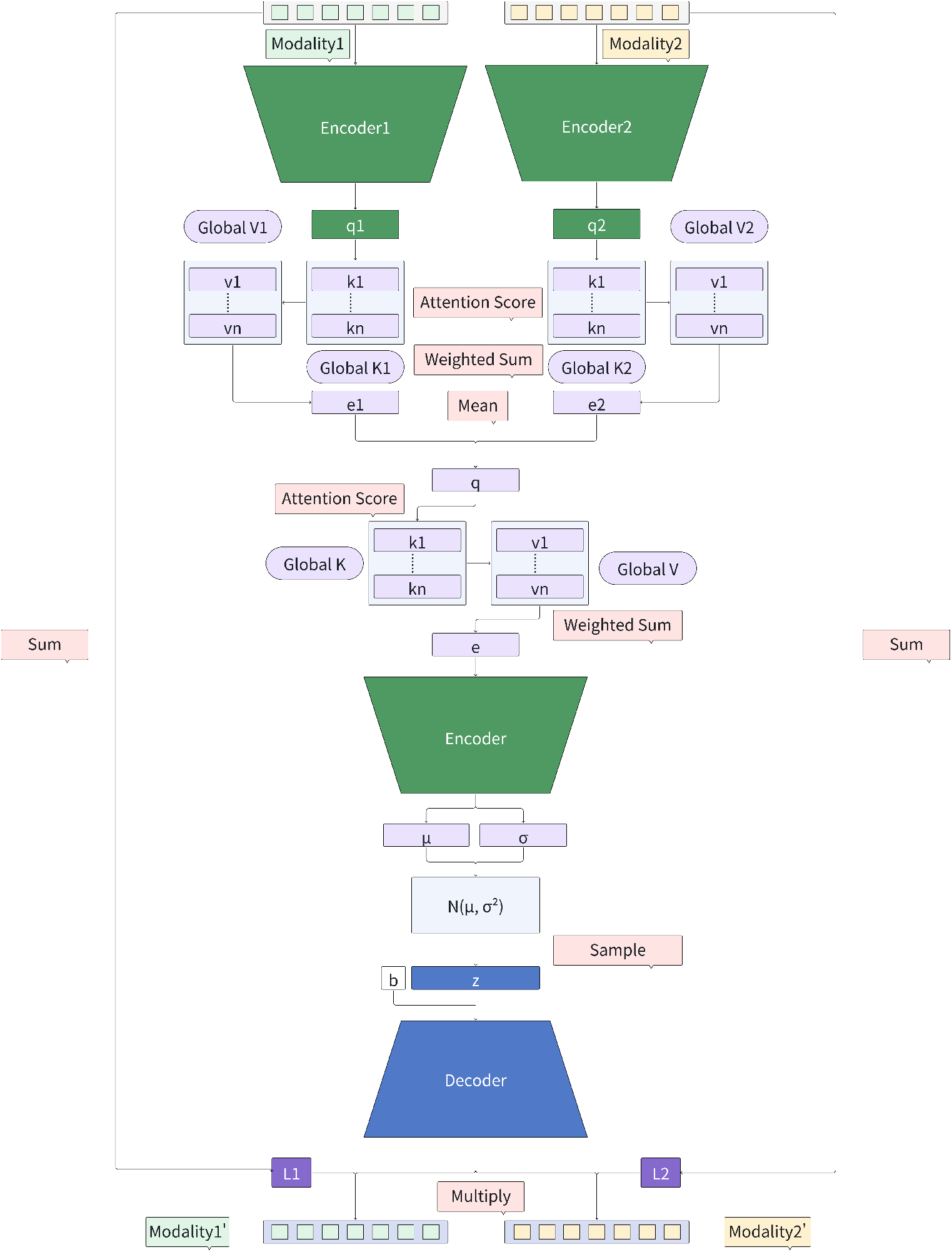
Overall architecture of the MultiGAI.

## 3 Results

To comprehensively evaluate the performance of MultiGAI, we conducted experiments on five datasets: two single-cell transcriptomics datasets, two single-cell multi-omics datasets, and one spatial transcriptomics dataset. For the single-cell transcriptomics datasets, the evaluation primarily focused on batch effects correction; for the single-cell multi-omics datasets, the evaluation tasks included latent variable quality, query mapping, and feature imputation; and for the spatial transcriptomics dataset, the evaluation focused on coordinate mapping and gene expression prediction.

The specific datasets used in this study are as follows:

- **Dataset 1** [17]: scRNA-seq data of mixed human and mouse immune cells, containing 97,861 cells, 8,135 genes, 23 batches, and 19 cell types.
- **Dataset 2** [17]: scRNA-seq data of human lung tissue, containing 32,472 cells, 15,148 genes, 16 batches, and 17 cell types.
- **Dataset 3** [18]: Multiome (scRNA-seq + scATAC-seq) data from bone marrow mononuclear cells (BMMCs) of healthy human donors. This dataset contains 69,249 cells, 13 batches, 13,431 genes, and 116,490 peaks, covering 22 cell types.
- **Dataset 4** [18]: CITE-seq (scRNA-seq + ADT) data from healthy human BMMCs. This dataset contains 90,261 cells, 12 batches, 13,953 genes, and 134 ADT features, covering 45 cell types.
- **Dataset 5** [19–28]: We integrated spatial transcriptomic data from 10 fresh-frozen mouse brain tissue slides and applied unified preprocessing to ensure a shared gene set across all slides. The resulting dataset contains 31,101 cells and 29,814 genes and is divided into 10 batches, with each slide treated as a separate batch.

In this section, MultiGAI and its variant adopt a unified model configuration. The random seed is fixed to 42 for all experiments, and the latent dimension is set to 30. The neural network architecture employs a single hidden-layer design, consisting of a linear layer, LayerNorm, ReLU activation, and Dropout, with a hidden width of 128. All models are trained for 200 epochs. Here, *α* denotes the weight of the inter-modality consistency loss based on cosine similarity, and *β* denotes the weight of the KL divergence loss. In all experiments, *α* is fixed to 1. During the first 100 epochs, *β* is set to 0 to avoid premature regularization, and during the remaining 100 epochs, *β* is set to 0.1 to gradually strengthen the regularization of the latent space. For single-cell transcriptomics and single-cell multi-omics tasks, the number of key–value pairs is set to 128, while for spatial transcriptomics tasks, it is set to 64. In addition, the weight of the spatial coordinate reconstruction loss, *λ*, is set to 100.

In the latent representation quality evaluation experiments, data preprocessing for GLUE [29] was strictly conducted according to its official tutorial; for all other methods, we followed the strategy of Multigrate [8] to select 4,000 highly variable genes and 20,000 highly variable peaks from the data. In the batch effects correction experiments, we likewise adopted the Multigrate [8] strategy to select 4,000 highly variable genes.

In the batch effects correction and latent variable quality evaluation experiments, we consistently employed the scIB framework [17] to assess model performance. This framework provides a variety of evaluation metrics. In this study, we selected six core metrics to comprehensively evaluate model performance from two perspectives: batch effects correction and biological information preservation (for detailed definitions and formulas of these metrics, see Appendix B). Moreover, in the batch effects correction experiments, we chose as baselines the methods that performed best in the scIB evaluation and directly compared them using the evaluation results already provided by scIB.

### 3.1 Batch Effect Correction

As comparative methods, we selected scVI [10], fastMNN [30], Scanorama [31], Seurat [6], and Harmony [32]. From the experimental results, MultiGAI achieved the highest overall scores in both datasets (see Table 1 and Fig. 2). Specifically, in Dataset1, MultiGAI achieved a biological preservation score close to the highest, while its batch effect correction score was substantially higher than those of other methods. In Dataset2, MultiGAI achieved scores close to the highest in both biological preservation and batch effect correction. Overall, MultiGAI effectively preserves biological information in single-cell transcriptomic data while achieving robust batch effect correction.

**Table 1.**
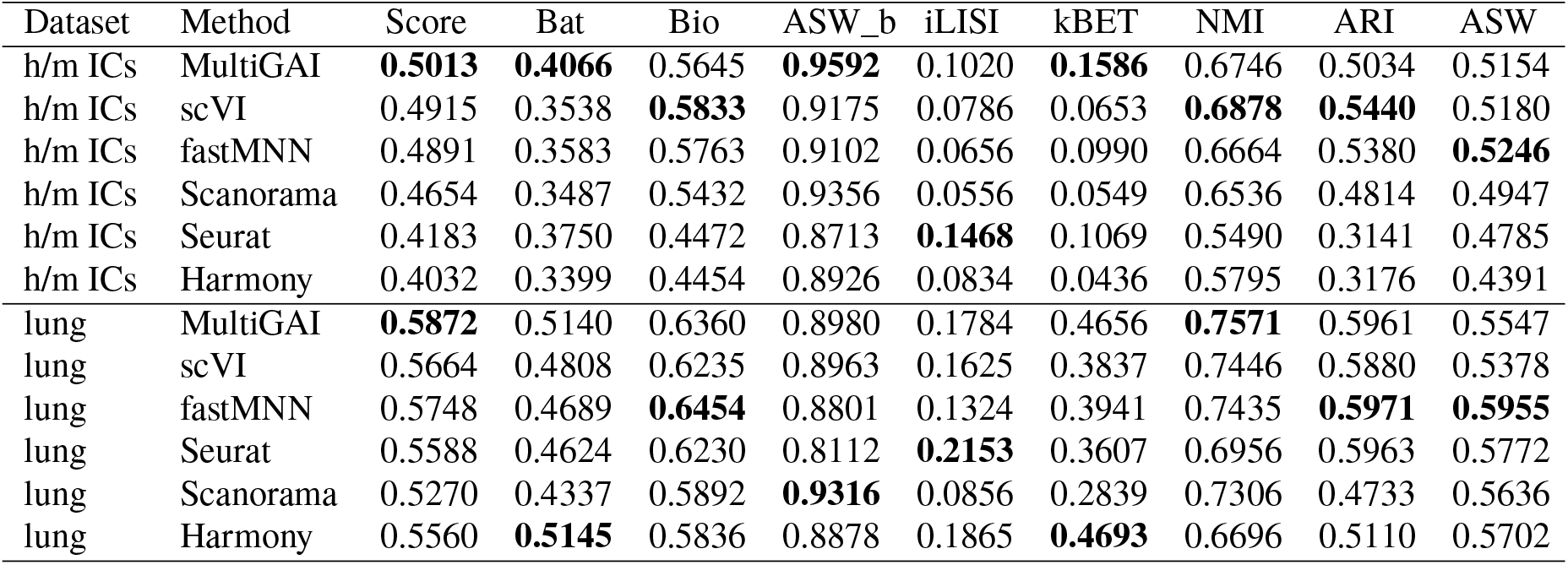
Metrics for h/m ICs and lung.

| Dataset | Method | Score | Bat | Bio | ASW_b | iLISI | kBET | NMI | ARI | ASW |
| --- | --- | --- | --- | --- | --- | --- | --- | --- | --- | --- |
| h/m ICs | MultiGAI | **0.5013** | **0.4066** | 0.5645 | **0.9592** | 0.1020 | **0.1586** | 0.6746 | 0.5034 | 0.5154 |
| h/m ICs | scVI | 0.4915 | 0.3538 | **0.5833** | 0.9175 | 0.0786 | 0.0653 | **0.6878** | **0.5440** | 0.5180 |
| h/m ICs | fastMNN | 0.4891 | 0.3583 | 0.5763 | 0.9102 | 0.0656 | 0.0990 | 0.6664 | 0.5380 | **0.5246** |
| h/m ICs | Scanorama | 0.4654 | 0.3487 | 0.5432 | 0.9356 | 0.0556 | 0.0549 | 0.6536 | 0.4814 | 0.4947 |
| h/m ICs | Seurat | 0.4183 | 0.3750 | 0.4472 | 0.8713 | **0.1468** | 0.1069 | 0.5490 | 0.3141 | 0.4785 |
| h/m ICs | Harmony | 0.4032 | 0.3399 | 0.4454 | 0.8926 | 0.0834 | 0.0436 | 0.5795 | 0.3176 | 0.4391 |
| lung | MultiGAI | **0.5872** | 0.5140 | 0.6360 | 0.8980 | 0.1784 | 0.4656 | **0.7571** | 0.5961 | 0.5547 |
| lung | scVI | 0.5664 | 0.4808 | 0.6235 | 0.8963 | 0.1625 | 0.3837 | 0.7446 | 0.5880 | 0.5378 |
| lung | fastMNN | 0.5748 | 0.4689 | **0.6454** | 0.8801 | 0.1324 | 0.3941 | 0.7435 | **0.5971** | **0.5955** |
| lung | Seurat | 0.5588 | 0.4624 | 0.6230 | 0.8112 | **0.2153** | 0.3607 | 0.6956 | 0.5963 | 0.5772 |
| lung | Scanorama | 0.5270 | 0.4337 | 0.5892 | **0.9316** | 0.0856 | 0.2839 | 0.7306 | 0.4733 | 0.5636 |
| lung | Harmony | 0.5560 | **0.5145** | 0.5836 | 0.8878 | 0.1865 | **0.4693** | 0.6696 | 0.5110 | 0.5702 |

**Figure 2.**
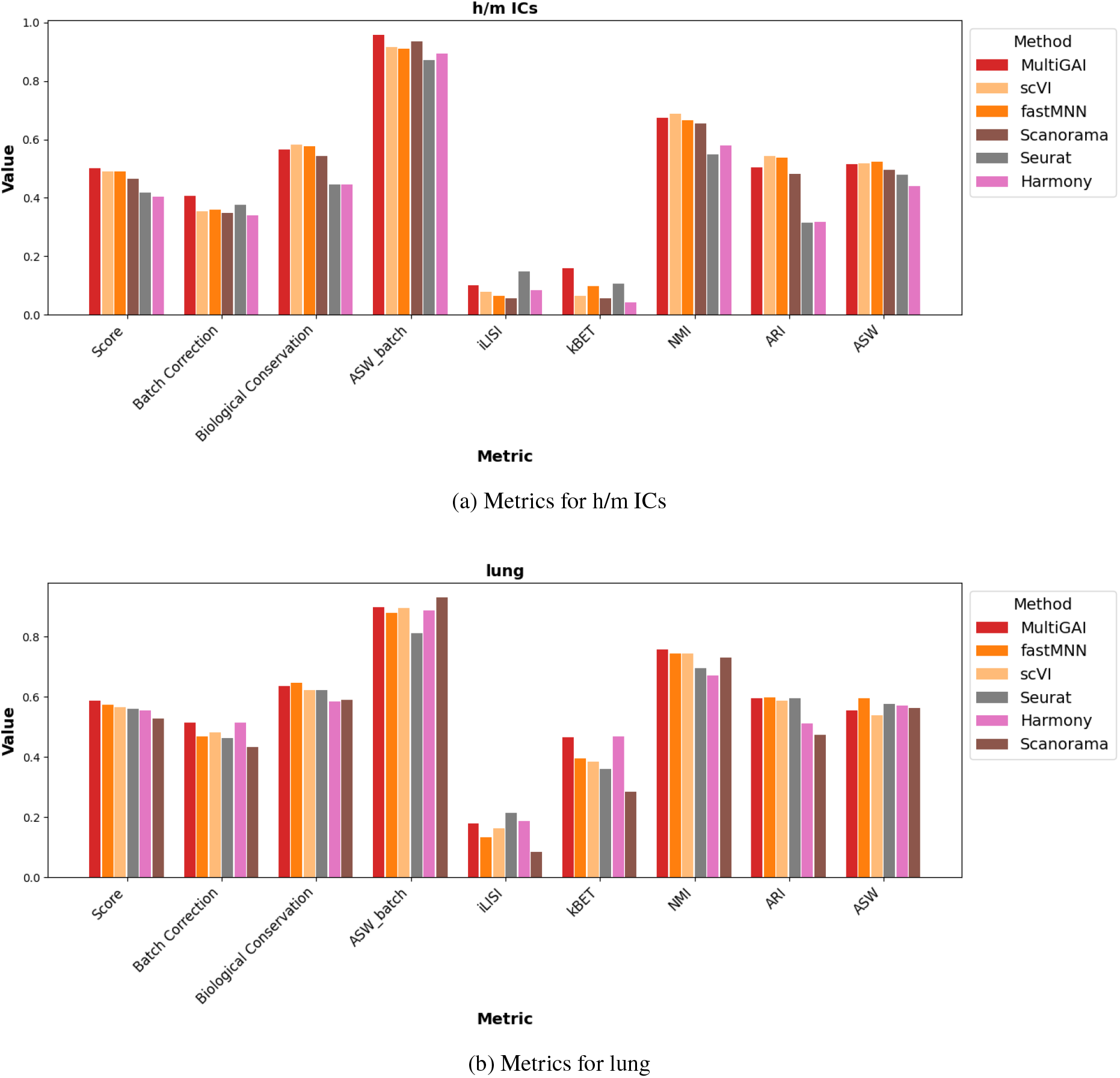
Metrics for h/m ICs and lung.

### 3.2 Latent Variable Quality

As comparative methods, we selected MultiVI [7] (and its CITE-seq variant totalVI [33]), GLUE [29], MOFA+ [34], MIMA [9], and Seurat [6]. For MIMA [9], the experimental setup followed the default configuration of the official tutorial, with the random seed fixed at 42 and the latent dimension set to 50. For GLUE [29], we evaluated each metric based on the latent representations of the integrated scRNA-seq and scATAC-seq data. For the remaining methods, aside from fixing the random seed at 42, all configurations followed the recommended settings of Multigrate [8].

Based on the experimental results, MultiGAI achieved the highest overall scores on both datasets, demonstrating its strong ability to correct batch effects while preserving biological variation (see Table 2 and Fig. 3). Specifically, in Dataset1, except for MIMA [9], MultiGAI achieved scores close to the highest in biological preservation while clearly outperforming other methods in batch effect correction. For MIMA [9], although it attained the highest batch effect correction score, its biological preservation score was slightly lower. In Dataset2, MultiGAI led in biological preservation scores while maintaining a relatively high level of batch effect correction. Overall, MultiGAI is able to generate high-quality latent representations in single-cell multiomics integration tasks under batch effect conditions.

**Table 2.** Latent variable quality metrics.

| Dataset | Method | Score | Bat | Bio | ASW_b | iLISI | kBET | NMI | ARI | ASW |
| --- | --- | --- | --- | --- | --- | --- | --- | --- | --- | --- |
| Multiome | MultiGAI | **0.6342** | 0.5777 | 0.6719 | 0.9201 | 0.3428 | 0.4703 | 0.7679 | **0.6965** | 0.5513 |
| Multiome | GLUE_ATAC | 0.5990 | 0.5102 | 0.6582 | 0.8854 | 0.3114 | 0.3339 | 0.7456 | 0.6534 | 0.5755 |
| Multiome | GLUE_RNA | 0.5960 | 0.4704 | **0.6797** | 0.8789 | 0.2836 | 0.2486 | **0.7808** | 0.6781 | 0.5801 |
| Multiome | MIMA | 0.6171 | **0.6018** | 0.6274 | **0.9663** | 0.3409 | **0.4980** | 0.7313 | 0.6378 | 0.5132 |
| Multiome | MOFA+ | 0.4392 | 0.2967 | 0.5343 | 0.8190 | 0.0246 | 0.0464 | 0.6424 | 0.4158 | 0.5447 |
| Multiome | MultiVI | 0.5897 | 0.4663 | 0.6719 | 0.8637 | 0.2827 | 0.2527 | 0.7730 | 0.6572 | **0.5856** |
| Multiome | Seurat | 0.5869 | 0.5068 | 0.6403 | 0.8051 | **0.3499** | 0.3655 | 0.7343 | 0.6013 | 0.5854 |
| CITE-seq | MultiGAI | **0.6273** | 0.5118 | **0.7044** | 0.9154 | 0.2914 | 0.3284 | **0.7946** | **0.7624** | 0.5562 |
| CITE-seq | MIMA | 0.5962 | 0.5015 | 0.6593 | **0.9524** | 0.2725 | 0.2798 | 0.7633 | 0.6883 | 0.5264 |
| CITE-seq | MOFA+ | 0.4206 | 0.2988 | 0.5018 | 0.8040 | 0.0135 | 0.0789 | 0.6328 | 0.3025 | 0.5701 |
| CITE-seq | Seurat | 0.5846 | **0.5352** | 0.6176 | 0.8014 | **0.3722** | **0.4319** | 0.7341 | 0.5288 | **0.5898** |
| CITE-seq | totalVI | 0.6001 | 0.5154 | 0.6566 | 0.9013 | 0.3063 | 0.3385 | 0.7865 | 0.6246 | 0.5586 |

**Figure 3.**
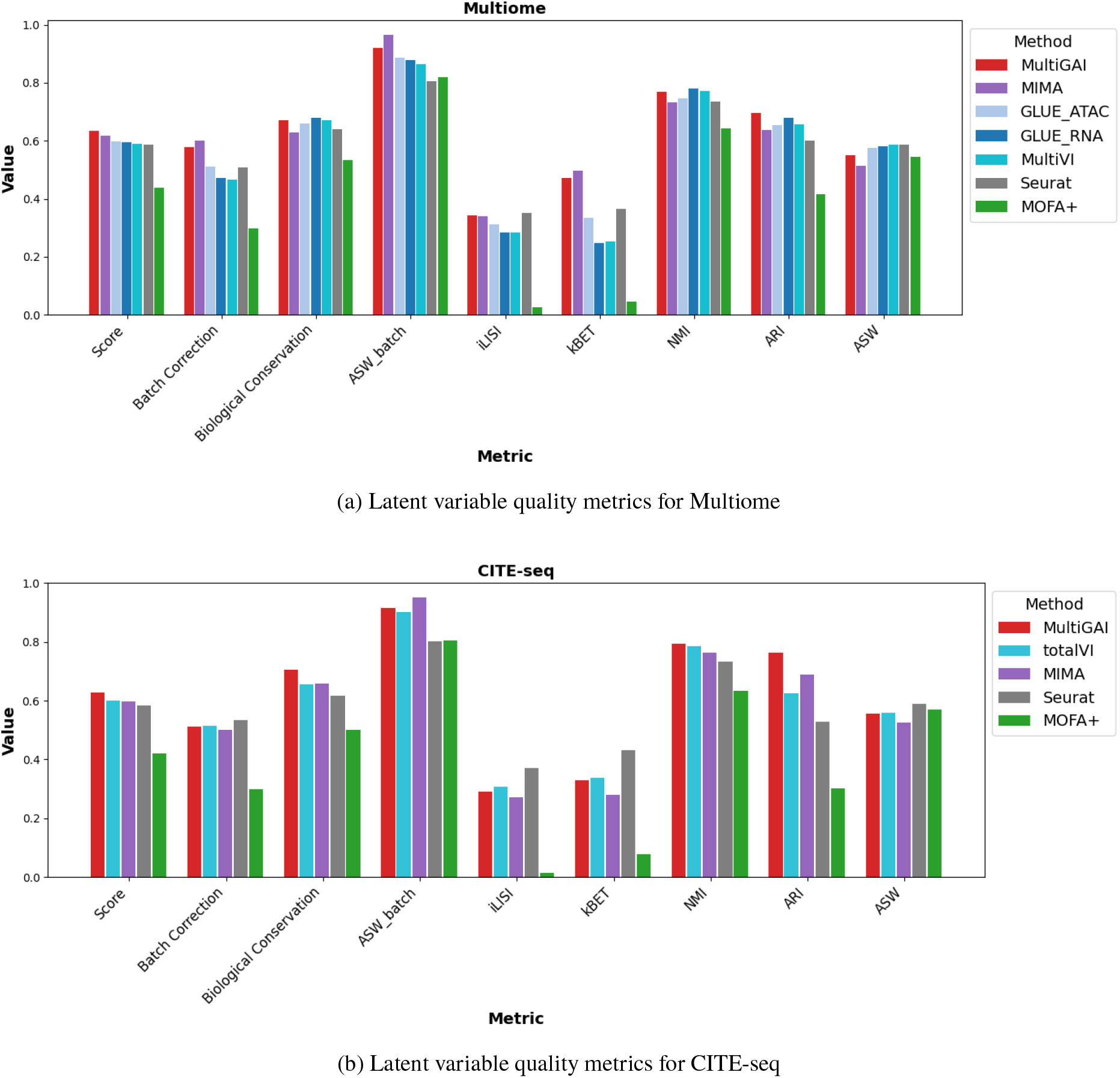
Latent variable quality metrics for Multiome and CITE-seq.

### 3.3 Query Mapping

To evaluate the query mapping capability of MultiGAI, we designed three experiments on each of two singlecell multi-omics datasets, and conducted two additional experiments on a combined dataset constructed by integrating these two datasets.

#### 3.3.1 Mapping of Single-Modality Data

For Dataset3 and Dataset4, we performed stratified sampling based on batch labels and cell type labels to obtain three subsets corresponding to data containing only scRNA-seq, only scATAC-seq (or only ADT), and both modalities, with relative proportions of 3:3:4, respectively. The data containing both modalities were used as the training set, and the trained model was then applied to infer the entire dataset. The UMAP [35] visualization was generated as follows: UMAP [35] parameters were first trained using the latent variables from the training set, and then the entire dataset was projected onto this UMAP [35] space. This experiment aims to evaluate the model’s ability to map data when only a single modality is provided.

Since the joint query vector in MultiGAI is the mean of the intermediate representations from all modalities, directly using the intermediate representation of each modality as the joint query vector is still inappropriate, even though modality consistency was enforced during training via a cosine similarity loss. Therefore, we added a simple neural network for each modality in MultiGAI to learn the mapping from that modality’s intermediate representation to the joint query vector. Specifically, after training the model on the training set, we collected the intermediate representations of each modality along with their corresponding joint query vectors, trained each neural network using mean squared error (MSE) loss, and then used these networks to map the intermediate representations of each modality to the joint query vectors.

From the UMAP visualizations [35] (see Figures 4 and 5), it can be observed that cells containing only a single modality are evenly mixed in the latent space with cells that have multi-omic data, without any noticeable structural shifts. These results indicate that MultiGAI is able to effectively map cells into a latent space consistent with multi-omic data, even when only a single modality is provided as input.

**Figure 4.**
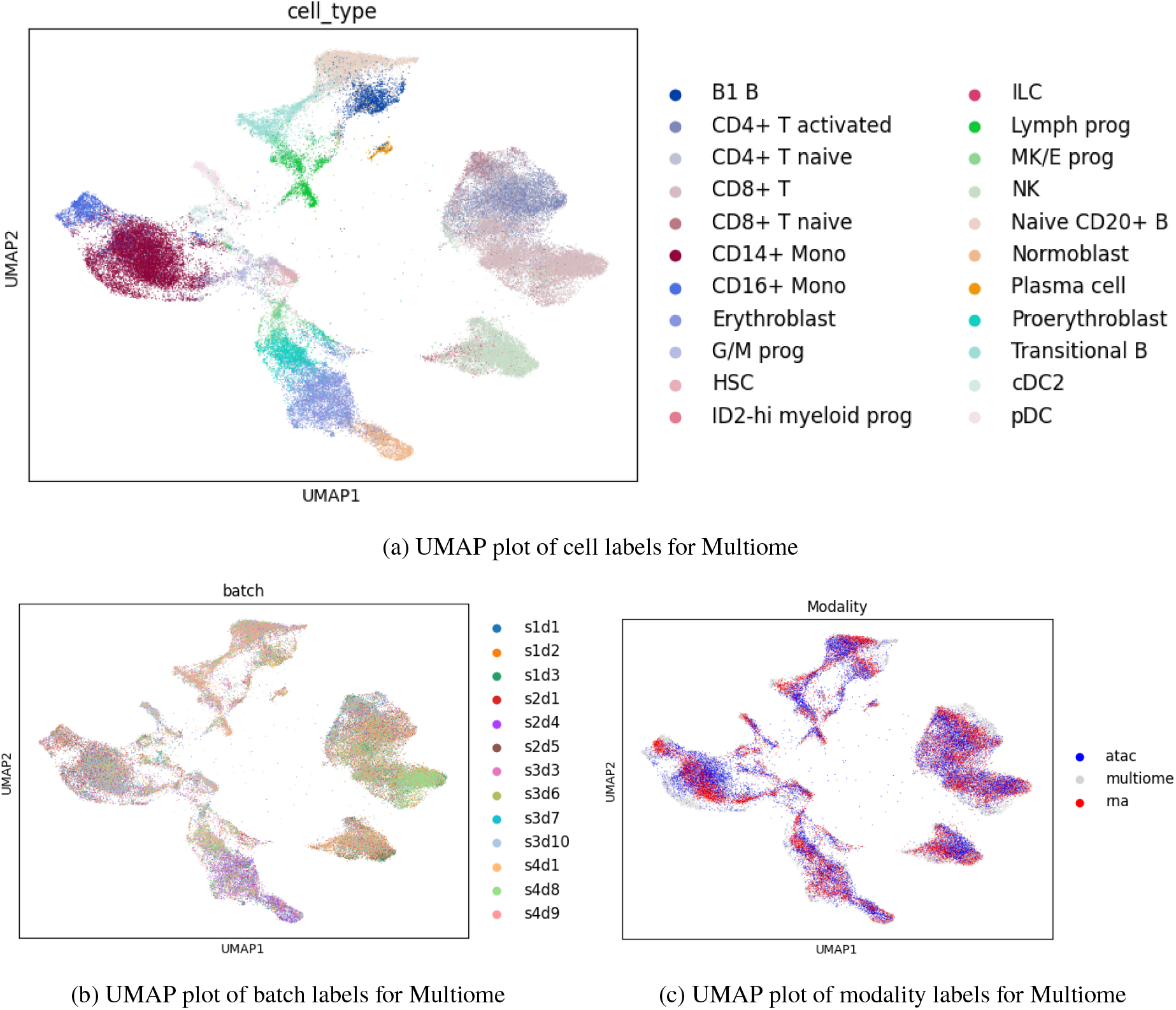
UMAP plots for Multiome dataset: cell labels (top), batch and modality labels (bottom).

**Figure 5.**
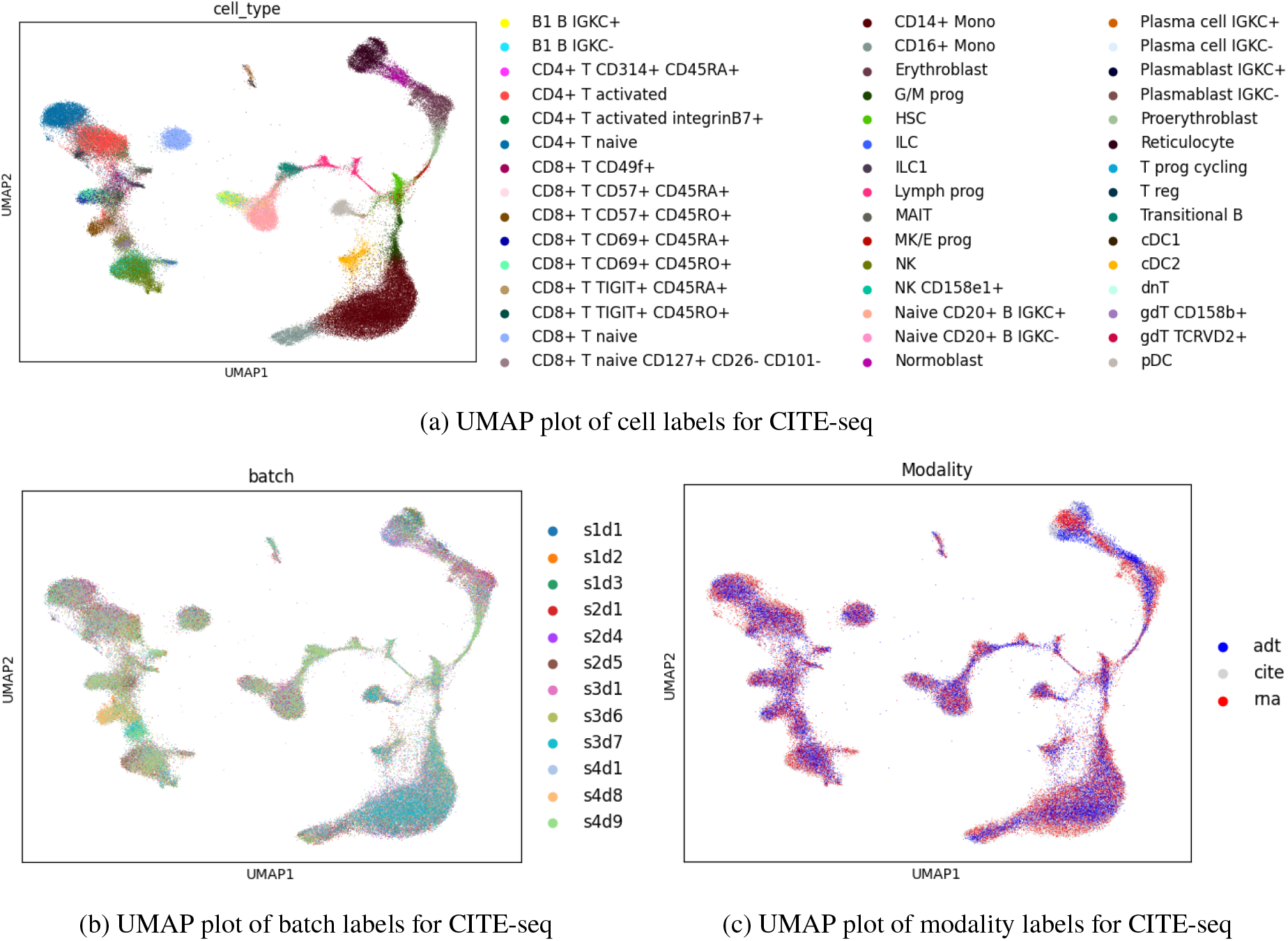
UMAP plots for CITE-seq dataset: cell labels (top), batch and modality labels (bottom).

#### 3.3.2 Mapping of Unseen Cell Types

For Dataset3 and Dataset4, we removed a cell type located at the edge of the UMAP [35] space and another located near the center, resulting in two datasets per original dataset, each lacking a specific cell type. These datasets with missing cell types were used as the training sets, and the trained models were then applied to infer the entire dataset. UMAP [35] visualizations were generated by projecting the latent variables of the full dataset. This experiment aims to evaluate the model’s ability to map unseen cell types.

From the UMAP [35] visualizations (see Figures 6 and 7), we observe that the held-out cell types do not mix with other distinct cell types; instead, they form separate clusters whose relative positions are largely consistent with those obtained using the full data. These results indicate that MultiGAI exhibits strong generalization ability and can correctly identify cell types that are similar to, but not present in, the training set.

**Figure 6.**
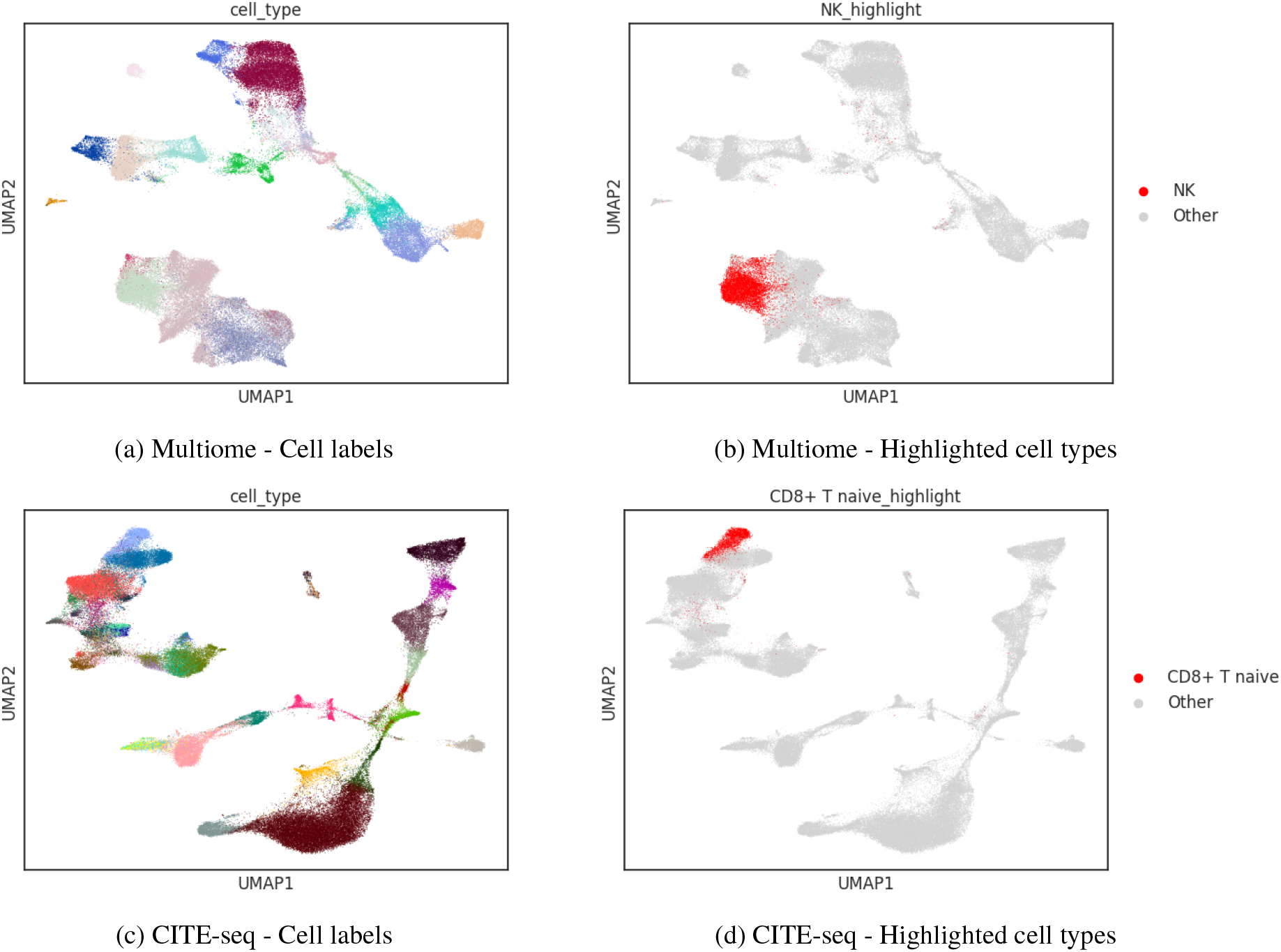
UMAP plots of edge cell labels and highlighted cell types for Multiome and CITE-seq datasets.

**Figure 7.**
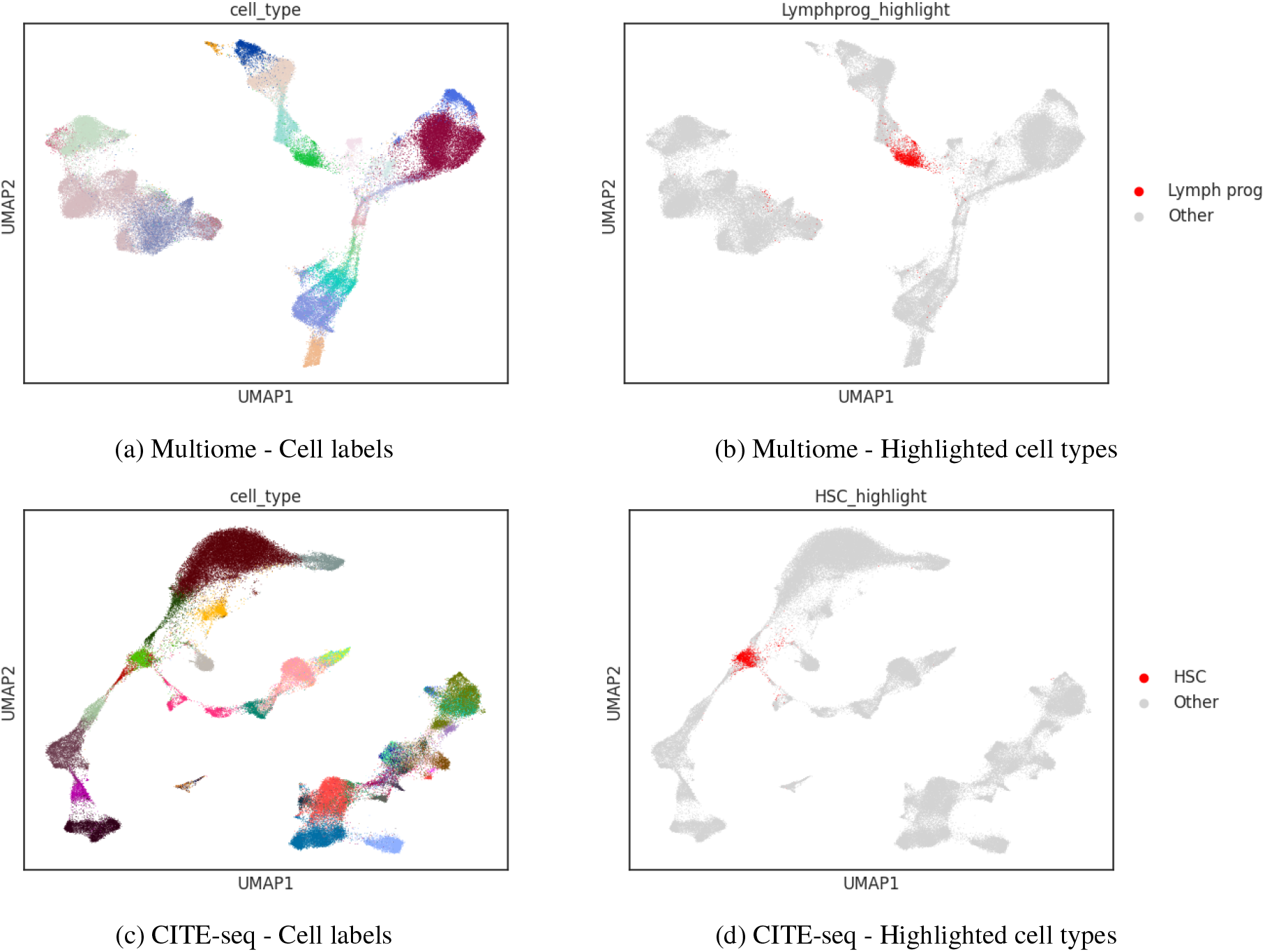
UMAP plots of center cell labels and highlighted cell types for Multiome and CITE-seq datasets.

#### 3.3.3 Integration and Mapping of Multiome and CITE-seq

To further evaluate the integration capability of MultiGAI, we first constructed a tri-modal dataset in which each sample contains only two modalities, based on the two datasets described above. Specifically, we first integrated the shared modality (scRNA-seq) from the two datasets, selected the common genes, and identified 4,000 highly variable genes using the same procedure as before. Next, for each sample, the missing modality was filled with zeros. This resulted in a tri-modal dataset containing 4,000 genes, 20,000 peaks (selected from Dataset3), and 134 ADT features. For different multi-omics datasets, we added modality-type labels to the existing batch labels in order to distinguish samples originating from different omics sources during integration.

After completing the integration experiment, we further performed a query mapping experiment. Following the stratified sampling strategy used in the *Mapping of Single-Modality Data* subsubsection, we partitioned the integrated dataset into subsets with a 1:1:1:2 ratio, corresponding to samples containing only scRNA-seq, only scATAC-seq, only ADT, and Multiome or CITE-seq, respectively. The proportion of Multiome and CITE-seq samples was kept consistent with their proportion in the integrated dataset. We then visualized the entire dataset using the same training procedure and UMAP [35] workflow as in the *Mapping of Single-Modality Data* subsubsection.

From the UMAP [35] visualizations (see Figure 8, where the top subfigures correspond to the integration task and the bottom subfigures correspond to the query mapping task), we observe that in both the integration and query mapping experiments, cells from different modalities are well mixed and cluster according to cell type. This demonstrates that MultiGAI possesses strong integration and mapping capabilities for single-cell multi-omics datasets that share at least one common modality.

**Figure 8.**
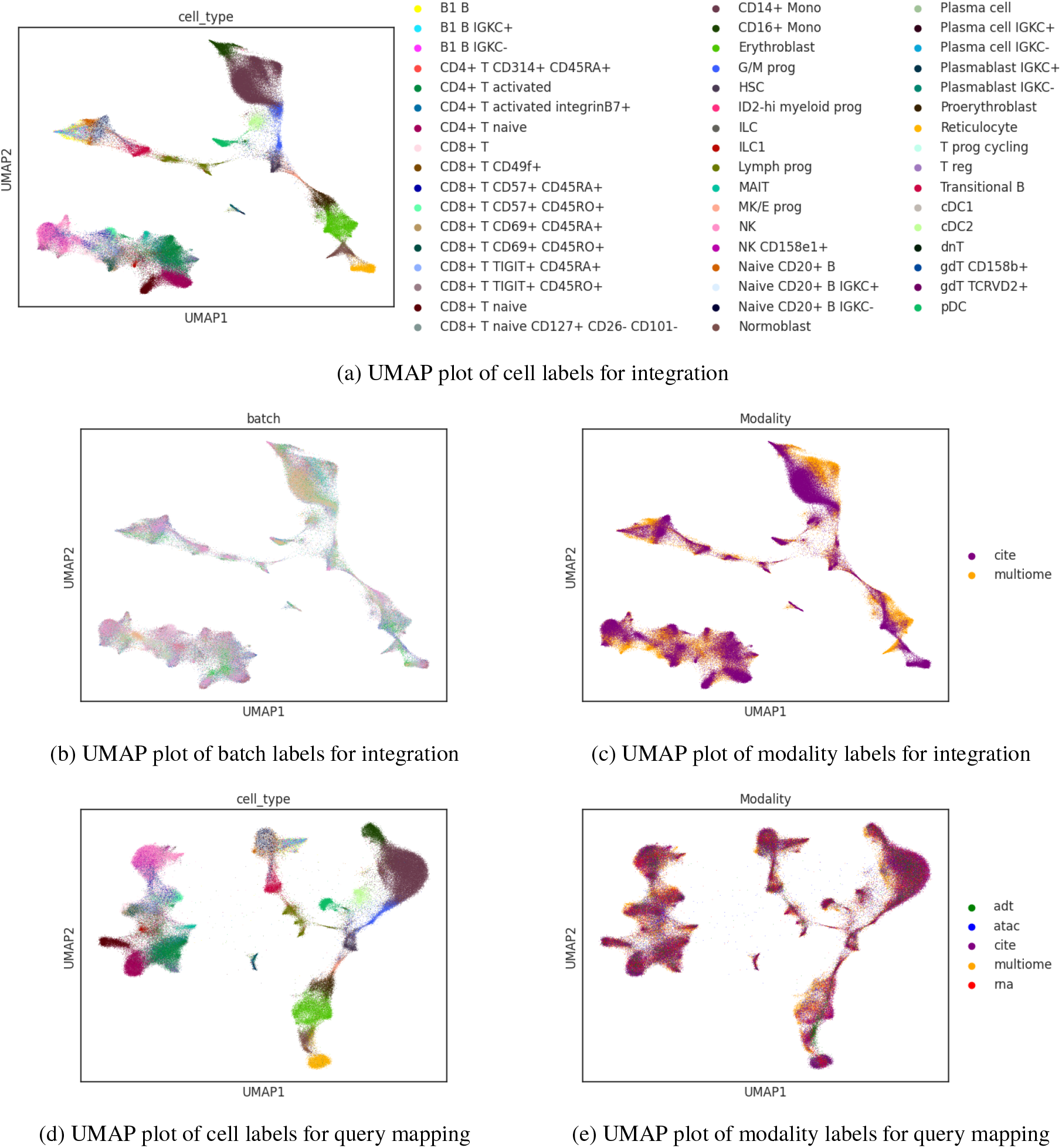
Integration and query mapping of Multiome and CITE-seq datasets.

### 3.4 Feature Imputation

To evaluate the feature imputation capability of MultiGAI, we conducted experiments on the integrated dataset used for query mapping. We selected several sets of canonical immune cell marker genes, their corresponding surface protein markers, and representative chromatin accessible regions to systematically assess MultiGAI’s imputation performance across scRNA-seq, ADT, and scATAC-seq modalities. Specifically, we focused on the following features:

- T cell-related features: the marker gene *CD3G*, the corresponding surface marker *CD3* [36], and a chromatin accessible region associated with *CD3G* expression, `chr11:118,343,914-118,344,801` [7];
- NK cell-related features: the marker gene *FCGR3A*, the corresponding surface marker *CD16*, which is primarily expressed in NK cells but can also be detected in CD16+ monocytes (non-classical/intermediate) [37];
- B cell-related features: the marker gene *MS4A1* and the corresponding surface marker *CD20* [38].

As shown in the figures (see Figures 9 and 10), the marker gene, surface protein, and chromatin accessible region for T cells all clearly delineate the T cell population; similarly, the marker gene and surface protein for B cells effectively identify the B cell population. For NK cells, both the marker gene *FCGR3A* and the surface marker *CD16* label NK cells as well as CD16+ monocytes, consistent with known biological observations [37].

**Figure 9.**
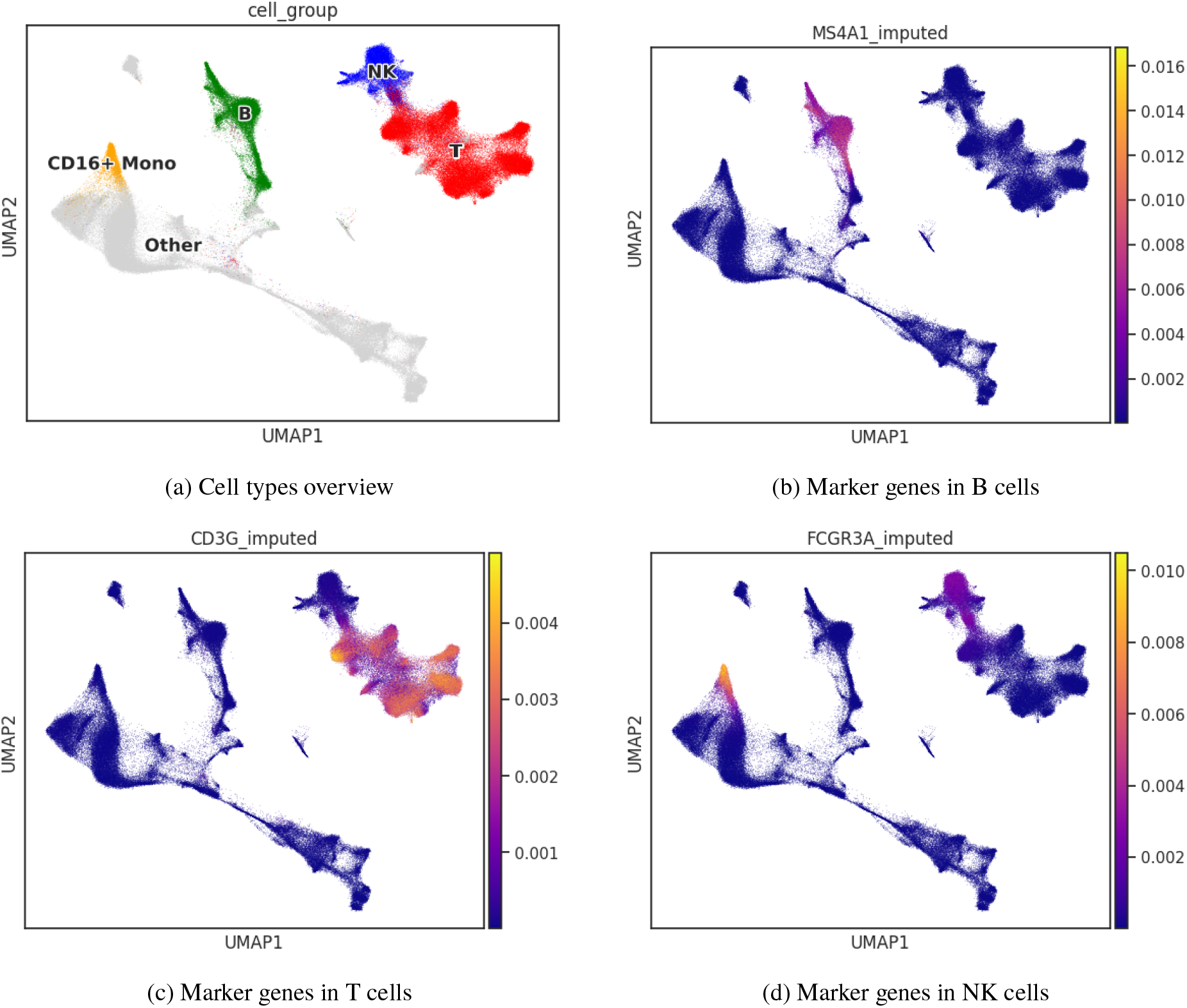
Marker genes across different cell types.

**Figure 10.**
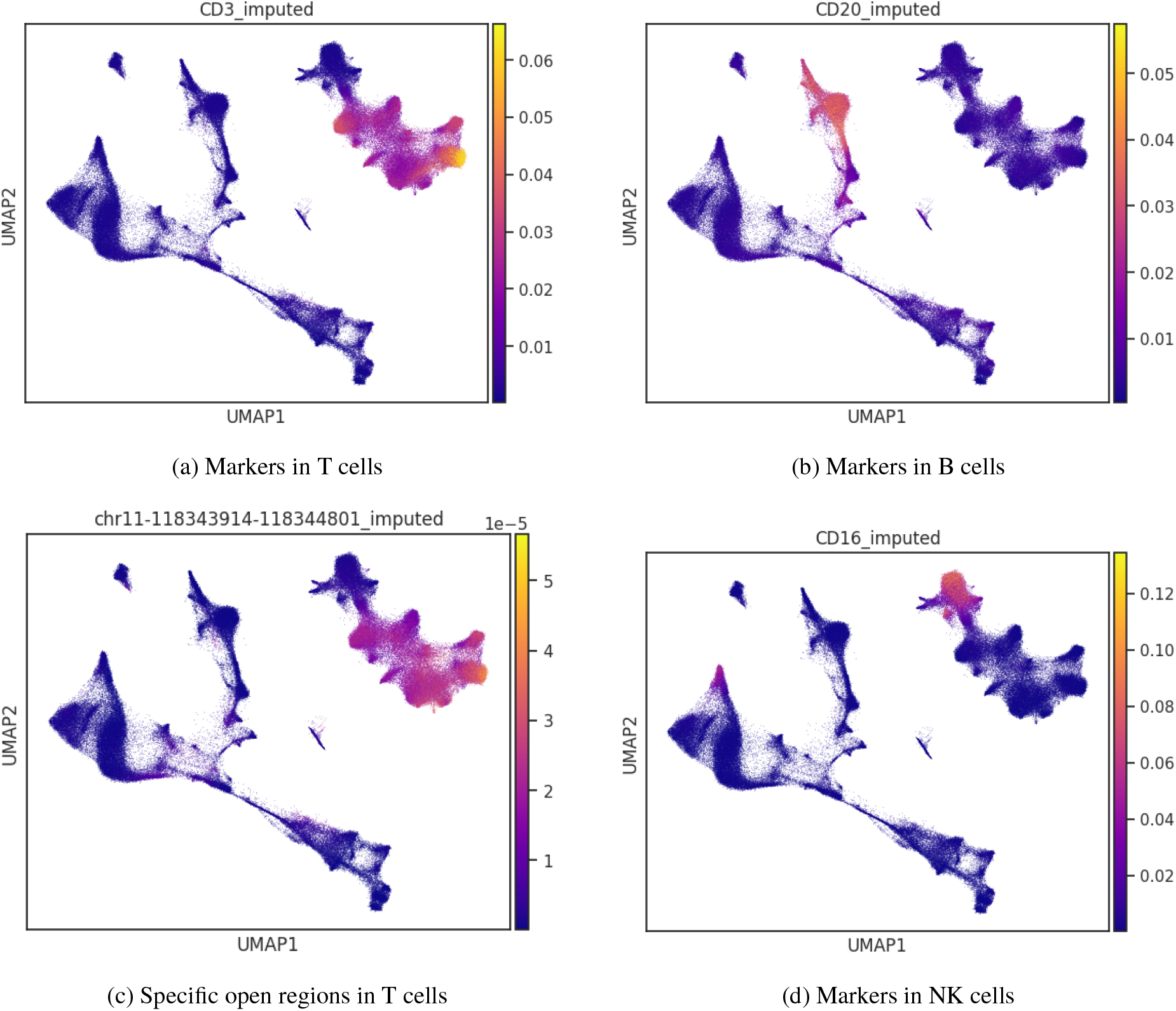
Markers and specific open regions across different cell types.

Notably, even when many cells contain only a single modality (scRNA-seq, scATAC-seq, or ADT), the gene expression, surface protein abundance, and chromatin accessibility imputed by MultiGAI exhibit consistent and stable patterns within the corresponding cell populations.

### 3.5 Evaluation of Spatial Transcriptomics Datasets

To evaluate the performance of MultiGAI on spatial transcriptomics data, we conducted two evaluation tasks on a fresh-frozen mouse brain spatial transcriptomics dataset: coordinate mapping and gene expression prediction.

#### 3.5.1 Coordinate Mapping

We performed stratified sampling by batch on the spatial transcriptomics dataset mentioned above, selecting 30% of the cells, which retained only the slide information and spatial coordinates. Then, we used the same training process as in the *Mapping of Single-Modality Data* subsection, applying the UMAP method [35] to visualize the entire dataset.

As shown in Figure 11, different batches (slides) are fairly evenly mixed together. However, it is worth noting that the latent variables of the spatial transcriptomics data are more scattered in the UMAP plot compared to single-cell multiomics data. This does not indicate that the model fails to integrate spatial transcriptomics data effectively, as each spatial transcriptomics sample typically includes multiple cells, and its gene expression reflects the combined result of these cells. Therefore, differences in gene expression patterns caused by variations in cell proportions are not batch effects but rather biological information. The model does not eliminate this biological variation, but rather causes the spatial transcriptomics data to appear more dispersed in the UMAP plot. For cells containing only spatial information, their latent representations are distributed uniformly within the population of cells whose latent representations comprise two ‘modalities’. The experimental results show that MultiGAI effectively integrates spatial transcriptomics data and maps cells containing only spatial information into the joint latent variable space.

**Figure 11.**
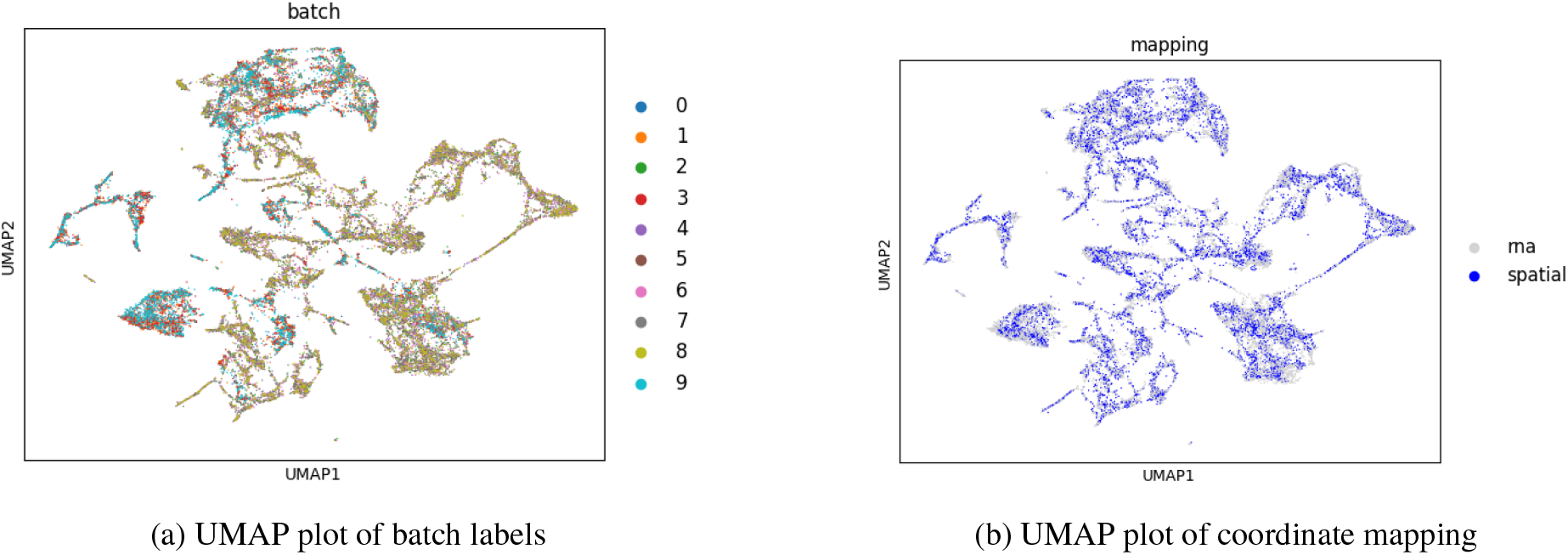
UMAP plot of spatial transcriptomics.

#### 3.5.2 Gene Expression Prediction

To evaluate the performance of MultiGAI in gene expression prediction, we selected several marker genes from mouse brain tissues, including *Cck, Mobp, Gng4*, and *Fabp7* [15, 39, 40]. In the coordinate mapping experiments, we selected a pair of sagittal sections from 10 slides, corresponding to the anterior and posterior regions. For each spatial coordinate point, we visualized the real expression levels and the predicted normalized expression levels of these genes, generating the corresponding spatial gene expression maps.

As shown in Figure 12, the predicted gene expression patterns are highly consistent with the real patterns, indicating strong performance in identifying and predicting marker genes. Because the model’s latent variables integrate spatial coordinate information, the predicted patterns exhibit clearer boundaries, smoother transitions, and more continuous expression compared to the real patterns, demonstrating that the model effectively captures the spatial continuity of gene expression. Furthermore, even for many cells that contain only spatial information, the model still achieves excellent prediction performance, reflecting its strong generalization ability.

**Figure 12.**
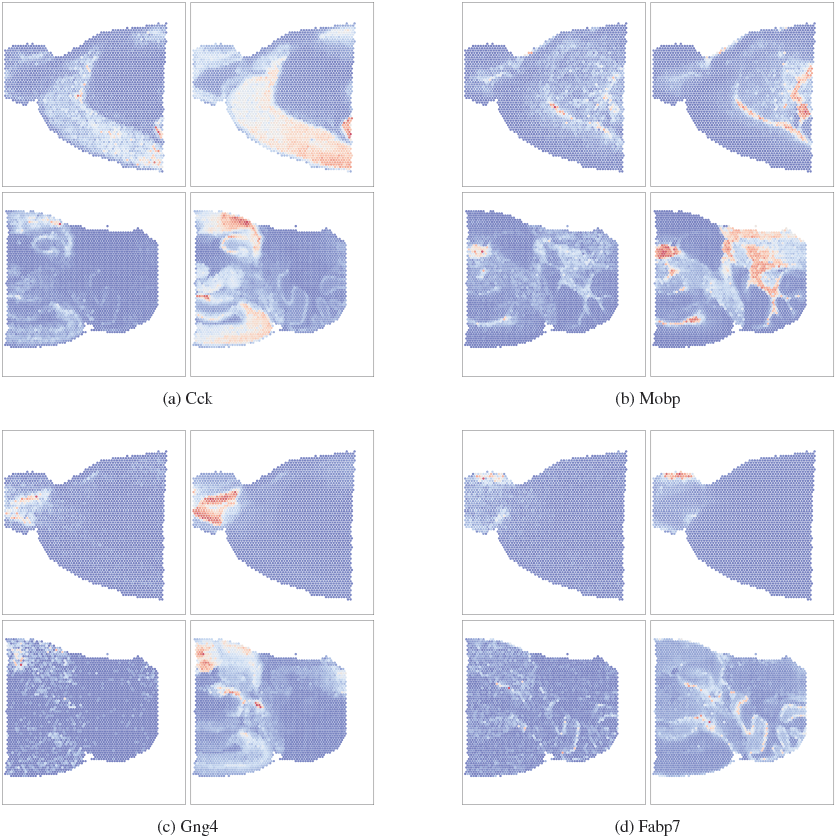
Spatial Gene Expression Maps for each gene (left: real expression, right: predicted normalized expression; top half: Anterior, bottom half: Posterior; top row: Cck and Mobp, bottom row: Gng4 and Fabp7)

### 3.6 Ablation Study

To evaluate the effectiveness of the global attention mechanism in MultiGAI, we designed an ablation model, MultiGAI_N, which has the same architecture and loss functions as MultiGAI but does not utilize the global attention mechanism. The metrics and datasets used in the ablation study are consistent with those in the latent variable quality experiments.

From the metric plots and tables (See Figure 13 and Table 3), we observe that while MultiGAI and MultiGAI_N achieve similar scores in biological conservation, MultiGAI significantly outperforms MultiGAI_N in batch effects correction. This indicates that incorporating the global attention mechanism can substantially improve batch effects correction while maintaining biological information, thereby demonstrating its effectiveness.

**Table 3.**
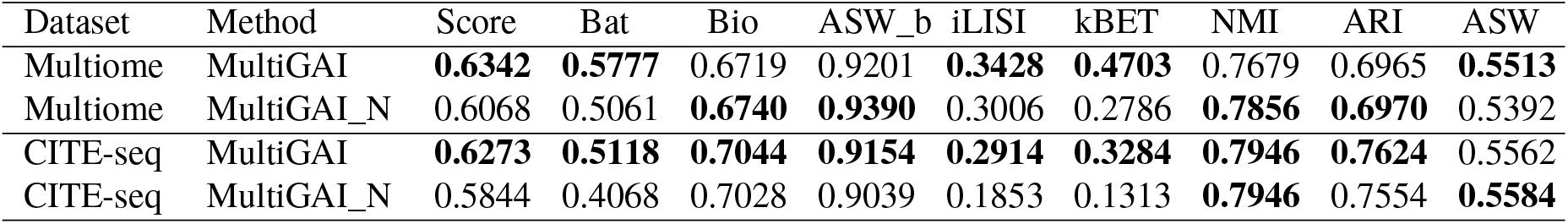
Ablation study metrics.

| Dataset | Method | Score | Bat | Bio | ASW_b | iLISI | kBET | NMI | ARI | ASW |
| --- | --- | --- | --- | --- | --- | --- | --- | --- | --- | --- |
| Multiome | MultiGAI | **0.6342** | **0.5777** | 0.6719 | 0.9201 | **0.3428** | **0.4703** | 0.7679 | 0.6965 | **0.5513** |
| Multiome | MultiGAI_N | 0.6068 | 0.5061 | **0.6740** | **0.9390** | 0.3006 | 0.2786 | **0.7856** | **0.6970** | 0.5392 |
| CITE-seq | MultiGAI | **0.6273** | **0.5118** | **0.7044** | **0.9154** | **0.2914** | **0.3284** | **0.7946** | **0.7624** | 0.5562 |
| CITE-seq | MultiGAI_N | 0.5844 | 0.4068 | 0.7028 | 0.9039 | 0.1853 | 0.1313 | **0.7946** | 0.7554 | **0.5584** |

**Figure 13.**
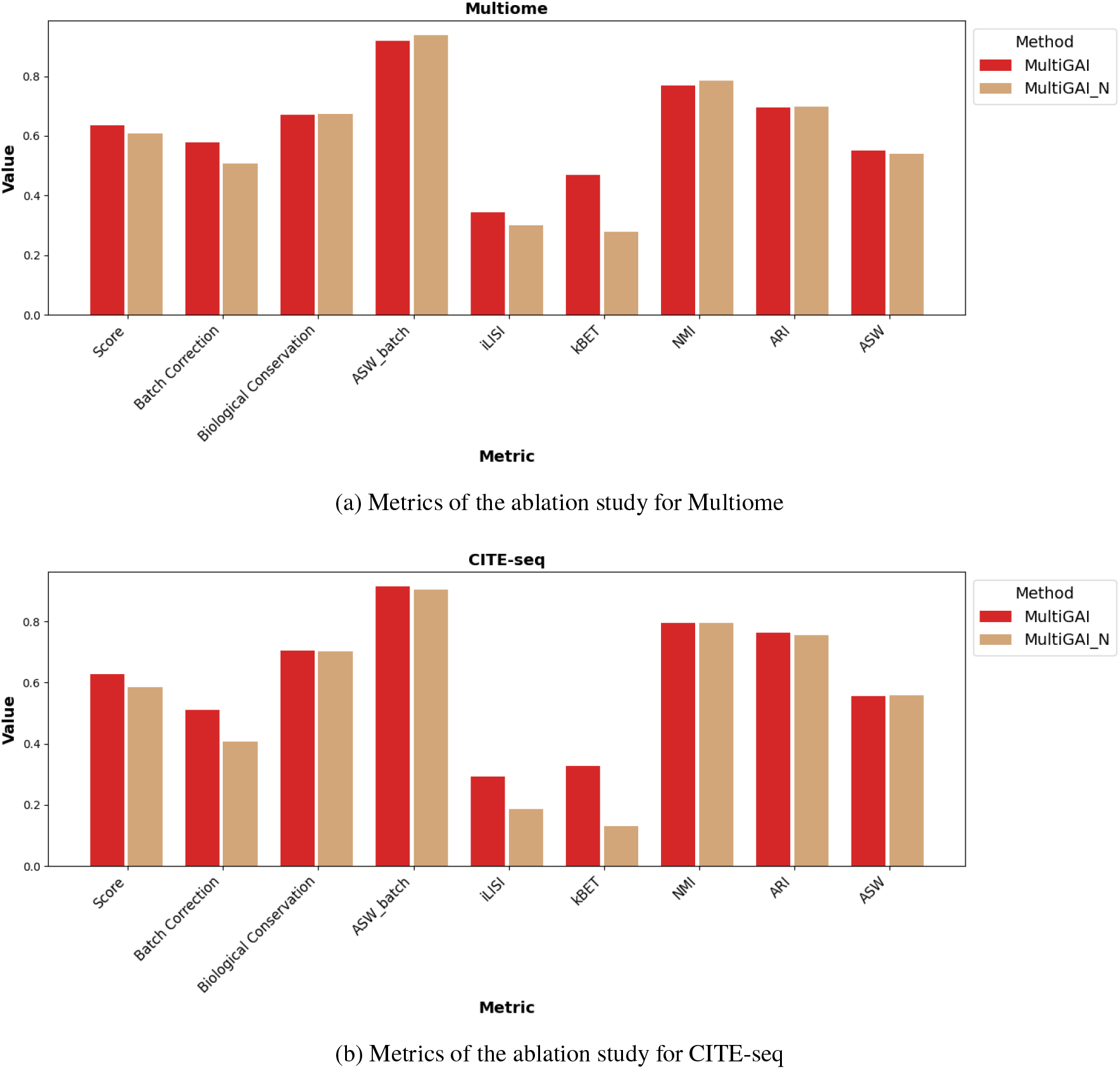
Metrics of the ablation studies for Multiome and CITE-seq datasets.

To evaluate the ability of the global attention mechanism to suppress batch effects while preserving the overall structure of the dataset, we extracted the joint query vectors, joint representations, and the set of joint global value vectors obtained from the integration experiments described in the *Integration and Mapping of Multiome and CITE-seq* subsection and performed visualization analysis. The UMAP plot (see Fig. 14) shows that the joint query vectors, when colored by cell type, still exhibit pronounced batch effects, whereas the joint representations processed by the global attention module effectively eliminate these batch effects, indicating that the global attention mechanism can effectively suppress the propagation of batch information. Furthermore, the UMAP plot (see Fig. 15) demonstrates that the joint global value vectors can effectively capture the overall structure of the dataset’s joint representations, suggesting that the global attention mechanism preserves the dataset’s global information.

**Figure 14.**
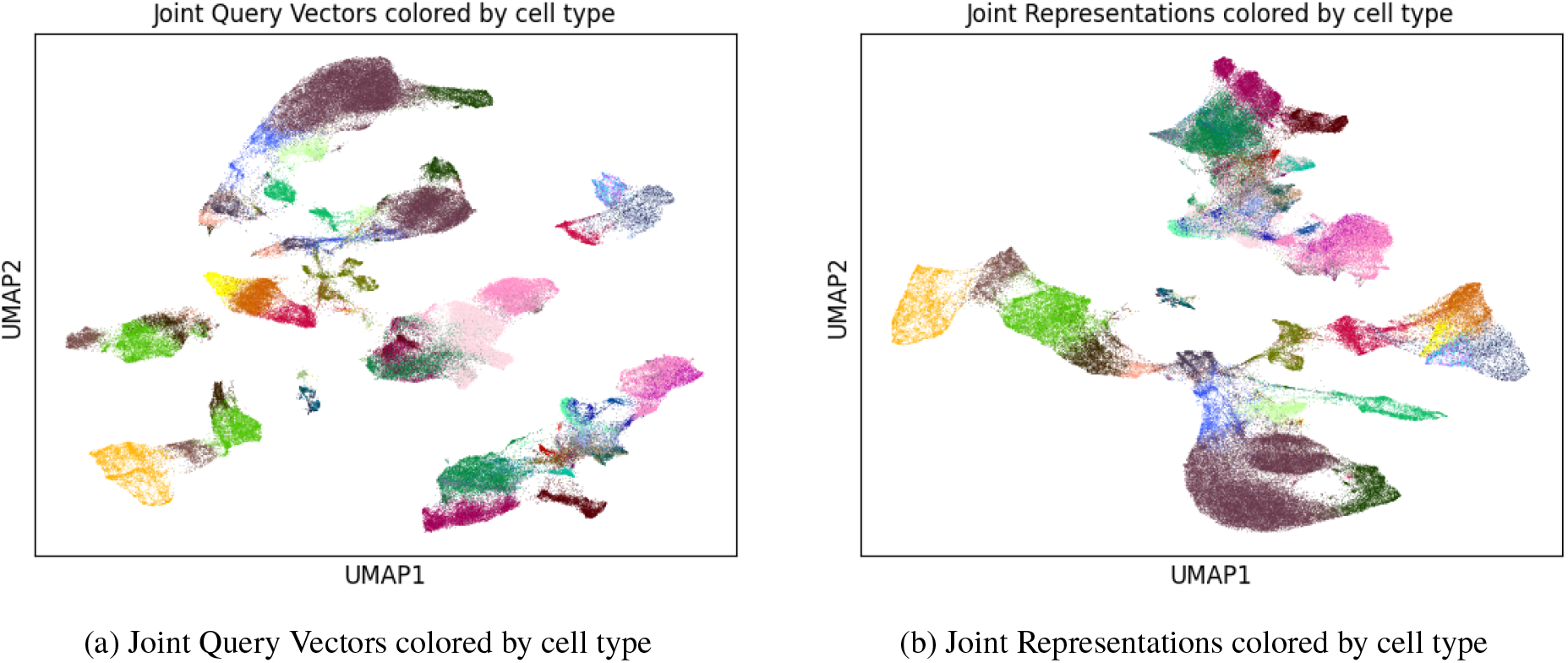
UMAP plots of joint query vectors and joint representations colored by cell type.

**Figure 15.**
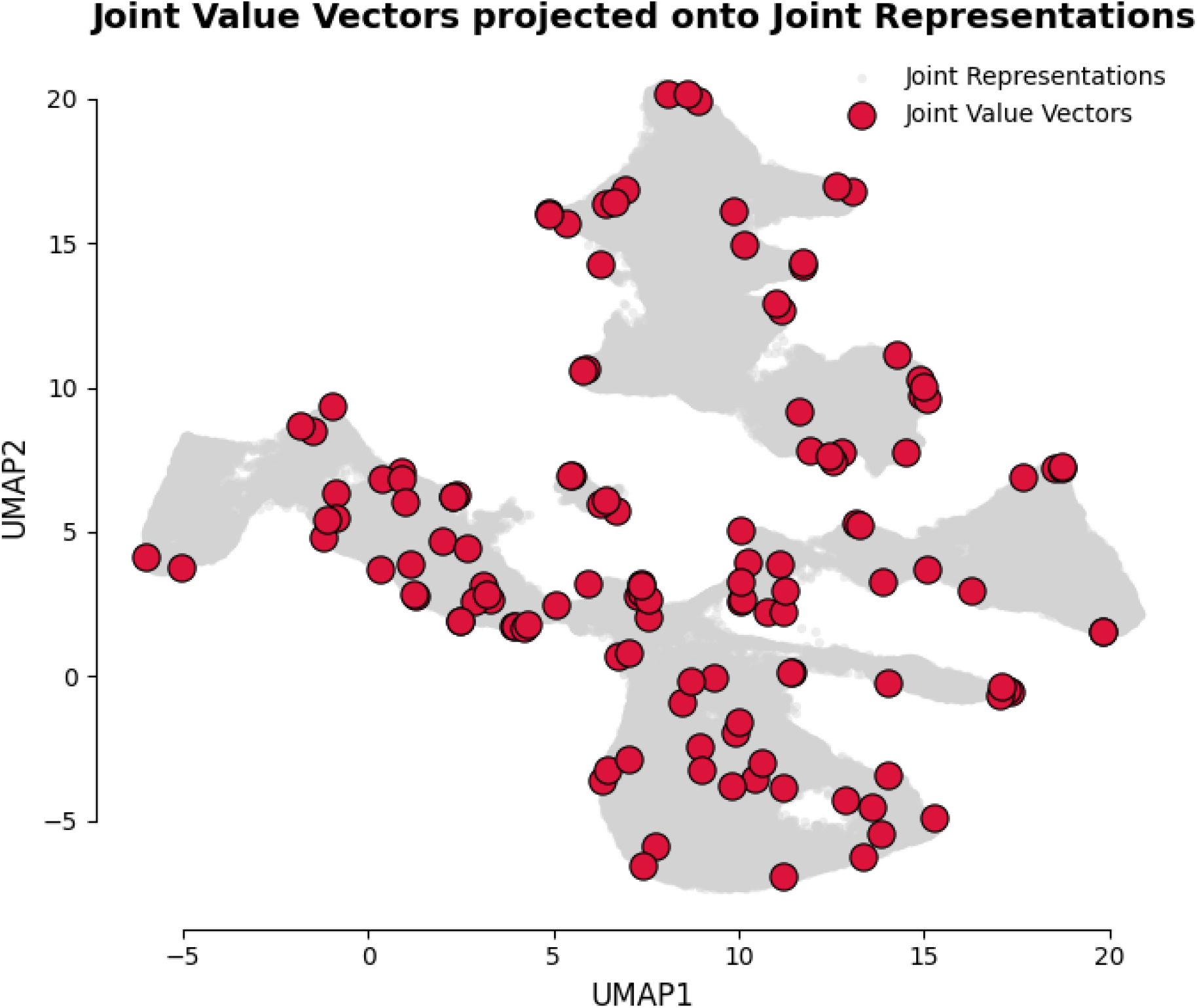
UMAP plots of joint value vectors and joint representations.

## 4 Conclusion and Outlook

To address the challenges faced by current single-cell multi-omics integration methods, this study proposes a variational autoencoder (VAE) framework with a global attention mechanism—**MultiGAI** [5]. MultiGAI can effectively remove batch effects while preserving biological information, thereby improving the integration of single-cell multi-omics data. The model not only exhibits strong integration capability and latent space mapping ability but also supports the integration of single-cell transcriptomic data and spatial transcriptomic data.

However, for spatial transcriptomic data, the current model primarily addresses the integration of gene expression data with spatial information, and has not yet tackled another challenge—**the deconvolution problem**. Therefore, one of the future research directions is to equip MultiGAI with deconvolution capability, further enhancing its applicability to spatial transcriptomic data. It should be noted that the decoder of MultiGAI mainly reconstructs the proportional structure of each modality rather than the raw count data. In future work, we plan to optimize the decoder to explore more accurate reconstruction strategies, thereby further expanding the model’s range of applications.

## Data Availability

All datasets used in this study are publicly available, including the single-cell multi-omics dataset (NCBI GEO, GSE194122) and fresh-frozen mouse brain spatial transcriptomics data (https://www.10xgenomics.com/). In addition, we used a human and mouse immune cell mixture dataset and a human lung tissue dataset [17], available at https://github.com/theislab/scib.

## Code Availability

The source code for this project is available on our GitHub: https://github.com/Recklesszjy/Multigai.

## A Detailed Model Formulation

### A.1 Normalization

Given spatial coordinates *x* and *y*, we normalize them to the range [0, 1] as follows:

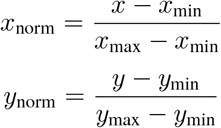

where *x*_min_, *x*_max_ and *y*_min_, *y*_max_ are the minimum and maximum values of the coordinates *x* and *y*, respectively.

### A.2 Fourier Transform Encoding

To enhance the spatial representation, we use multi-frequency Fourier positional encoding. For a given coordinate **c**, the encoding is:

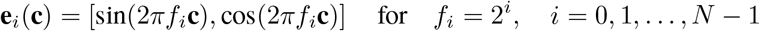

where *f*_*i*_ represents the frequency and *N* is the number of frequency levels.

### A.3 Encoder

The encoder maps single-cell multi-omics or spatial transcriptomics data into a unified latent space using a global attention mechanism. For a single input sample with multiple modalities 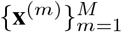, where *M* denotes the number of modalities, the encoder first projects each modality (except the spatial modality) into a query vector in the hidden dimension:

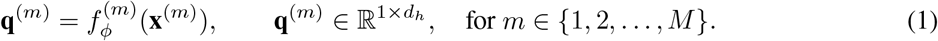

For sequencing data (denoted as *m* = *R*), the input **x**^(*R*)^ consists of the raw count data:

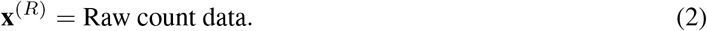

For each modality (except the spatial modality), a trainable key-value set is defined in the hidden dimension:

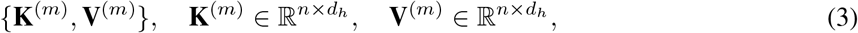

where *n* is the number of key-value pairs and *d*_*h*_ is the hidden dimension.

The modality-specific intermediate representation is computed via attention:

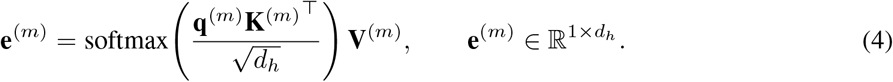

For the spatial modality of spatial transcriptomics data (denoted as *m* = *S*), the input **x**^(*S*)^ consists of the concatenation of the one-hot encoded slide ID and the spatial coordinates, which are first normalized and then encoded using Fourier transform encoding:

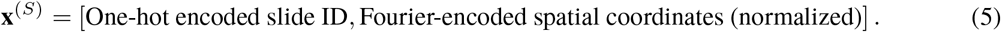

The intermediate representation of the spatial modality is obtained via a neural network:

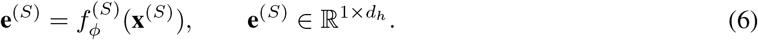

Intermediate representations from all modalities are averaged to obtain a joint query vector:

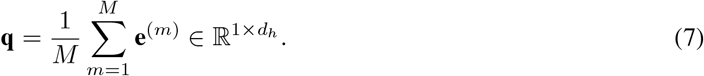

A joint global key-value set is defined in the hidden dimension:

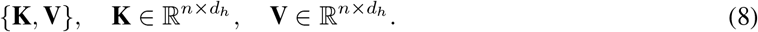

The unified representation is computed via attention between the joint query vector **q** and the joint key-value set:

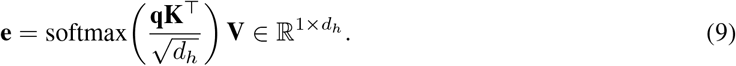

Finally, the joint representation **e** is fed into two separate linear layers to predict the mean and standard deviation of the latent variable:

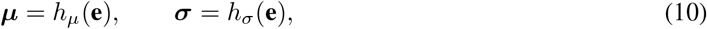

where 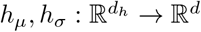 and *d* is the latent dimension.

### A.4 Latent Variable Sampling

We employ the reparameterization trick to sample latent variables from a Gaussian distribution:

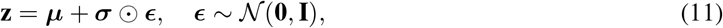

where **z** ∈ ℝ^1×*d*^ denotes the sampled latent vector.

### A.5 Decoder

The decoder takes as input the concatenation of the latent variable **z** and the batch label **b**:

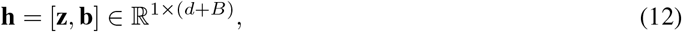

where **b** ∈ {0, 1} ^*B*^ is a one-hot encoding of the batch label and *B* is the total number of batches.

For sequencing data (denoted as *m* = *R*), the decoder predicts the parameters of a zero-inflated negative binomial (ZINB) distribution:

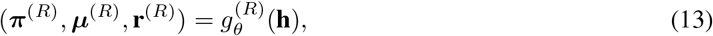

where ***µ***^(*R*)^ is normalized via softmax to represent relative expression proportions.

The sequencing depth is given by:

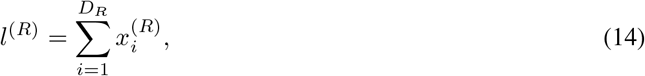

and the expected expression is scaled as:

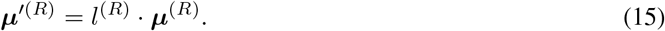

For spatial data (denoted as *m* = *S*), the decoder directly predicts the normalized spatial coordinates **c**^(*S*)^:

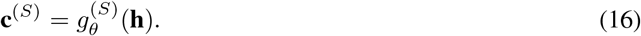

### A.6 Zero-Inflated Negative Binomial (ZINB) Distribution

For a single feature *x*_*i*_ in sequencing data, the ZINB distribution is defined as a mixture of a point mass at zero and a Negative Binomial (NB) distribution:

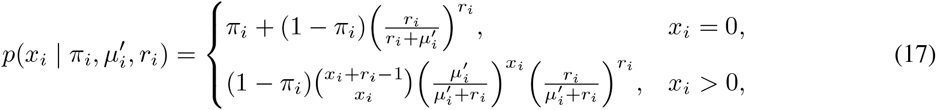

where *π*_*i*_ ∈ [0, 1] is the zero-inflation probability, 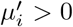 is the mean of the NB component, and *r*_*i*_ *>* 0 is the dispersion parameter.

### A.7 Loss Function

The training objective minimizes a composite loss:

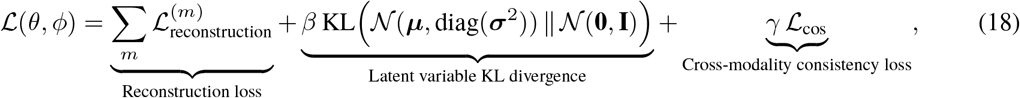

where **h** = [**z, b**], and *β, γ >* 0.

1. Reconstruction loss (ℒ_**reconstruction**_) The modality-specific reconstruction losses are defined as follows:
  - **Sequencing data (***m* = *R*): the negative expected log-likelihood under the predicted ZINB distribution

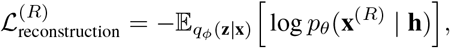

where **x**^(*R*)^ denotes the observed sequencing counts, and *p*_*θ*_(**x**^(*R*)^ | **h**) is modeled by a ZINB distribution with parameters *π, µ*^′^, *r*.
  - **Spatial data (***m* = *S*): weighted mean squared error (MSE) for normalized spatial coordinates

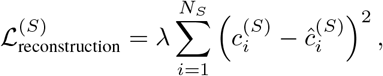

where *N*_*S*_ is the number of coordinates, 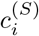 is the true normalized coordinate, 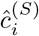 is the predicted coordinate, and *λ >* 0 is the weight.
2. Cross-modality consistency loss (ℒ _cos_) This term enforces similarity between intermediate representations of different modalities. Denoting 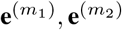 as the modality-specific intermediate representations for two modalities, the loss is defined as:

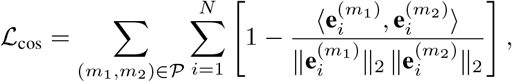

where *N* is the number of samples, 𝒫 is the set of all modality pairs, 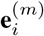 is the intermediate representation of sample *i* in modality *m*, and ⟨·, ·⟩ denotes the inner product.

## B Metrics

To comprehensively evaluate the quality of the latent variables learned by MultiGAI, we employed the scIB [17] framework for integration quality assessment. We selected six core metrics covering two key dimensions:

- **Batch-effect Correction Metrics** (weight: 40%):

**–** `Batch ASW`: Batch mixing silhouette width
**–** `iLISI`: Local inverse Simpson’s index for batch mixing
**–** `kBET`: k-nearest neighbor batch effect test

- **Biological Conservation Metrics** (weight: 60%):

**–** `Cell-type ASW`: Cell-type silhouette width
**–** `NMI`: Normalized mutual information (between clustering and true labels)
**–** `ARI`: Adjusted Rand index (between clustering and true labels)

The overall score is calculated as:

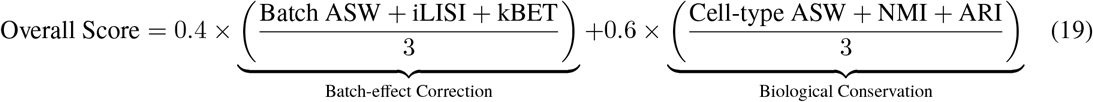

